# Metric Learning on Expression Data for Gene Function Prediction

**DOI:** 10.1101/651042

**Authors:** Stavros Makrodimitris, Marcel J.T. Reinders, Roeland C.H.J. van Ham

**Affiliations:** Delft Bioinformatics Lab, Delft University of Technology, Van Mourik Broekmanweg 6, 2628XE, Delft, the Netherlands; Keygene N.V., Agro Business Park 90, 6708PW, Wageningen, the Netherlands; Leiden Computational Biology Center, Leiden University Medical Center, Einthovenweg 20, 2333ZC, Leiden, the Netherlands

## Abstract

**Motivation:** Co-expression of two genes across different conditions is indicative of their involvement in the same biological process. However, using RNA-Seq datasets with many experimental conditions from diverse sources introduces batch effects and other artefacts that might obscure the real co-expression signal. Moreover, only a subset of experimental conditions is expected to be relevant for finding genes related to a particular Gene Ontology (GO) term. Therefore, we hypothesize that when the purpose is to find similar functioning genes that the co-expression of genes should not be determined on all samples but only on those samples informative for the GO term of interest.

**Results:** To address both types of effects, we developed MLC (Metric Learning for Co-expression), a fast algorithm that assigns a GO-term-specific weight to each expression sample. The goal is to obtain a weighted co-expression measure that is more suitable than the unweighted Pearson correlation for applying Guilt-By-Association-based function predictions. More specifically, if two genes are annotated with a given GO term, MLC tries to maximize their weighted co-expression, and, in addition, if one of them is not annotated with that term, the weighted co-expression is minimized. Our experiments on publicly available *Arabidopsis thaliana* RNA-Seq data demonstrate that MLC outperforms standard Pearson correlation in term-centric performance.

**Availability:** MLC is available as a Python package at www.github.com/stamakro/MLC

**Supplementary information:** Supplementary data are available online.

## 1 Introduction

Knowing which biological processes and pathways are affected by each gene would be a useful tool for plant biologists and breeders. With this information, they can more easily identify genes that are likely to affect the phenomenon or trait they are studying and prioritize genes for experimental testing. The Biological Process Ontology (BPO) of the Gene Ontology (GO) (Ashburner *et al*., 2000) provides us with a set of terms that describe biological processes at different levels of granularity and can be used to annotate genes from all species in a systematic way. However, the use of computational methods to accurately predict BPO annotations, also known as Automatic Function Prediction (AFP), remains challenging, as demonstrated in the Critical Assessment of Functional Annotation (CAFA) challenges (Jiang *et al*., 2016a).

Most AFP methods use the Guilt-By-Association (GBA) principle. They define a similarity or dissimilarity measure between genes and use it as a proxy for functional similarity. Then, they assign GO annotations to genes of unknown function based on the functions of the genes most similar to them. The choice of similarity measure is always motivated by biology. For instance, sequence similarity points towards a conserved structure which in turn implies similar function. Alternatively, co-expression across different conditions may hint at involvement in the same pathways. Combining multiple similarity measures in order to better approximate functional similarity is also possible, as done for instance in (Lan *et al*., 2013; Cozzetto *et al*., 2013; Zhang *et al*., 2017).

Genes that are involved in the same biological processes are expected to show similar expression patterns, as they respond similarly to perturbations related to these processes. Discovering BPO annotations for all unannotated genes requires data from a wide range of different experimental conditions. For example, we need samples from different tissues, different time points across development, from wild-type or mutant plants etc. Thanks to world-wide sequencing efforts, more and more RNA-Seq data are becoming available to public databases, such as ArrayExpress (Parkinson *et al*., 2007) and GEO (Clough and Barrett, 2016).

The Pearson Correlation Co-efficient (*PCC*) is the most widely used measure of gene co-expression similarity and has been largely successful, especially for microarray-derived expression data. For instance, for MS-kNN (Lan *et al*., 2013), one of the top-performing methods in CAFA2 (Jiang *et al*., 2016a), the PCC was calculated on samples from 392 human microarray datasets to quantify co-expression similarity between genes, outperforming sequence similarity for AFP in BPO (Lan *et al*., 2013).

PCC might, however, not be the optimal co-expression measure due to the diversity of biological processes and heterogeneity of public expression datasets. Firstly, only a subset of all available experimental conditions is likely to be truly informative about a specific GO term. For example, let us assume that we are looking for genes involved in plant immune response. Using the PCC across all possible conditions, we implicitly expect that all such genes are expressed similarly not only during immune response, but across all conditions and tissues. However, differential co-expression analysis has shown that several *Arabidopsis thaliana* immune genes, such as FLS2, ADR1 and JAR1, change co-expressed partners before and after infection with Pseudomonas syringae (Jiang *et al*., 2016b). A gene that is co-expressed with immune genes during (only) infection is still a good candidate gene for immune response, even if it has different expression patterns to the immune genes in other tissues or developmental stages. Including many unrelated expression samples, essentially adds noise to the correlations. Secondly, batch effects and other technical biases that are not corrected during pre-processing can introduce noise and corrupt the co-expression signal. PCC treats all samples equally and implicitly assumes that there are no systematic differences between them. According to these reasonings, we should be able to improve the performance of coexpression-based gene function prediction by calculating co-expression only over the samples that are relevant for each term.

This insight that the PCC might be suboptimal is not new. For example, Jaskowiak et al. showed that k-means clustering of gene expression data heavily relies on the choice of similarity measure (PCC, Spearman correlation, Euclidean distance etc.) and that the most suitable measure varies across different datasets (Jaskowiak *et al*., 2012). As another example, Hu et al. showed that using an inappropriate distance metric can really harm the performance of the *k*-Nearest Neighbors (*k*-NN) classifier in biomedical datasets (Hu *et al*., 2016).

Adapting a distance measure is a subfield within machine learning that is called metric learning: learning a distance function from a dataset of examples that can most effectively be utilized to perform a task, e.g. discriminating between two classes. It is most explored in combination with the *k*-NN classifier (Bellet *et al*., 2013). In the context of AFP, Ray and Misra developed a metric learning method called Genetic Algorithm for Assigning Weights to Gene Expressions using Functional Annotations (*GAAWGEFA*) that learns a weighted PCC on microarray data using a genetic algorithm to find the optimal values for the weights (Ray and Misra, 2019). They showed that their weighted correlation increases the protein-centric precision compared to PCC in a yeast dataset. Metric learning has also been applied to AFP combined with multiple-instance learning (Xu *et al*., 2017). In that work, each protein is viewed as a “bag of domains” and metric learning is used to learn a distance function between proteins (based on their domains) that is representative of functional similarity. Here, we use metric learning to identify the most informative conditions for a given GO term. Similar to GAAWGEFA, our goal is to assign a weight to every RNA-Seq sample. GAAWGEFA learns one weighted PCC for all GO terms (Ray and Misra, 2019). On the contrary, our approach, Metric Learning for Co-expresssion (MLC), optimizes the weights per term. Our philosophy (graphically shown in Figure 1) is that weights should be chosen in such a way that a pair of genes annotated with the same term should have maximally similar expression profiles, i.e. comply with our assumption that these genes should be co-expressed. On the other hand, when one gene is annotated with the term and the other not, we expect that such a pair should not have high co-expression. In other words, we would like to select weights that minimize the co-expression for these pairs. For pairs of genes both not annotated with the term, we cannot say anything about the co-expression since they might be annotated with another term, and thus be co-expressed too (albeit for other conditions/samples). Consequently, the co-expression of these pairs should be ignored when optimizing weights for the GO term of consideration. A high weight for a sample will put emphasis on that sample when calculating the co-expression over all samples, whereas a low weight for a sample will reduce the influence of that sample. When a weight becomes zero the sample is even ignored. To enforce selecting informative samples, we additionally apply an L1 sparsity constraint on the weights, which will set a weight to zero when a sample is uninformative (Tibshirani, 1996). In contrast to GAAWGEFA, where they have used a genetic algorithm to find the weights, we are able to pose the weight optimization in an elegant mathematical formulation that can be minimized efficiently using standard methods. To reduce the computational burden even further, we use the weighted inner product as a similarity function instead of the weighted PCC. We evaluate our algorithm on public RNA-Seq data from *A. thaliana*.

**Figure 1.**
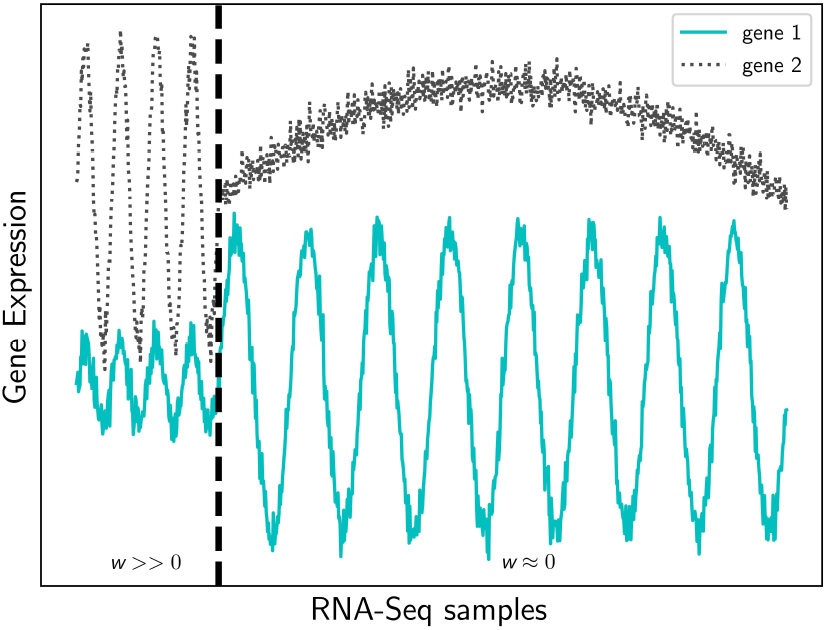
Illustrative example of the expression of two hypothetical genes (*y-*axis, solid and dashed lines) involved in the same biological process over a large set of samples (*x*-axis). The total Pearson correlation between the genes is 0.09. MLC sets large weight values (*w*_*m*_) for the samples left of the vertical dashed line (where the unweighted correlation is 0.92) and small or zero weights for the samples on the right (unweighted correlation = 0.002).

## 2 Methods

### 2.1 Data and Preprocessing

We used the API of the European Bioinformatics Institute (EBI) (Petryszak *et al*., 2017) to download all *A. thaliana* RNA-seq studies available at ArrayExpress (Parkinson *et al*., 2007). All samples had been processed using the same pipeline and expression was measured using raw read counts. We restricted our dataset to samples that used the latest version of the *A. thaliana* genome (TAIR10) and had fewer than 10% unmapped reads. After removing duplicate experiments, we had 4,215 samples from 298 different studies (batches) for 32,833 genes, 26,925 of which were protein-coding. We used a preprocessing pipeline similar to the one used to construct the ATTED-II RNA-Seq co-expression network (Obayashi *et al*., 2018). We first removed samples with fewer than 10,000,000 mapped reads. Then, we removed lowly expressed genes (genes with maximum expression over the remaining samples less than 100). To diminish the zero-inflation of the dataset, we also removed all genes that were not expressed in at least half of the samples (median expression smaller than 1). We then mapped the TAIR gene ID’s to UniProt ID’s. BPO annotations were downloaded from GOA (https://www.ebi.ac.uk/GOA) in September 2016 and annotations with the IEA evidence code were removed. A total of 2,978 samples and 6,013 genes with BPO annotations remained after these filtering steps. We applied ComBat (Johnson *et al*., 2007) to remove unwanted variation stemming from the fact that the different samples come from different studies (batch effects). ComBat uses a Bayesian method to standardize the mean and the variance of each gene in each study (batch). In order to be able to estimate within-batch variances, we removed all studies that had only one sample, leaving us with 2,959 samples.

### 2.2 Notation

We use **x**_*i*_ ∈ ℝ^*f*^ to denote the expression of gene *i* across all *f* = 2,959 samples. *x*_*im*_ is the expression of gene *i* at sample *m*. 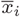 is the mean of gene i across all samples. Given *N* genes and a GO term *l*, we denote as y(*l*) ∈ [0, 1]^*N*^ the vector with the class labels of the genes, with **y**(*l*)_*i*_ = 1 iff gene *i* is annotated with *l*. The sample weights are represented by a vector with *f* non-negative elements **w**(*l*) ∈ [0, +∞)^*f*^.

### 2.3 Weighted and Unweighted Measures of Co-Expression

The most widely-used measure of co-expression between two gene expression vectors **x**_*i*_, **x**_*j*_ is the Pearson Correlation Coefficient (*PCC*), which is defined as follows:

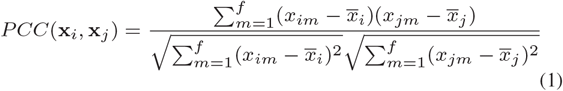

Note that the numerator is the covariance between **x**_*i*_ and **x**_*j*_ and the denominator is the product of the standard deviations of the two vectors.

A related, but simpler measure is the inner product similarity (*S*), which, on the contrary, is sensitive to the mean expression of both genes.:

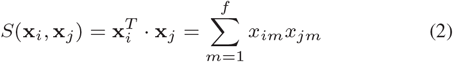

If two vectors **x**_*i*_, **x**_*j*_ both have zero mean and unit L2-norm, then their *PCC* is equal to their inner product. This equality does not hold anymore if we weigh each vector element (sample) differently. However, since the two metrics are related, we chose to use the weighted inner product similarity instead of the weighted PCC as our expression similarity function in order to simplify the problem. We center and scale our data so that the (unweighted) mean of every gene is zero across all conditions and its (unweighted) L2-norm is equal to one:

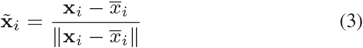

Then, we define our similarity function as the weighted inner product of the two scaled expression vectors (*S***_w_**):

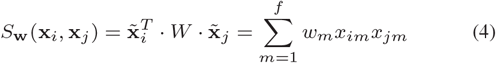

Where *W* = *diag*(**w**) is a diagonal matrix containing the sample weights.

### 2.4 Metric Learning for Co-expression (MLC)

The rationale for learning the weights is to maximize the performance of the *k*-NN classifier. For this purpose, we want the expression similarity between two genes that are both annotated with a given GO term *l* to be higher (on average) than the similarity between a gene that is annotated with *l* and a gene that is not. We group each gene pair into one of the following three categories:

1. both genes are annotated with *l* (we call these “positive-positive pairs” or “*p*-*p*”),
2. exactly one of the two genes is annotated with *l* (“positive-negative pairs” or “*p*-*n*”) and
3. neither gene is annotated with *l* ((“negative-negative pairs” or “*n*-*n*”)).

Our goal is to find the weight values *w*_*m*_ that maximize the separability between “*p*-*p*” and “*p*-*n*” pairs. Let *μ*_*p*−*p*_, 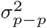 denote the mean and variance of the weighted similarity value *S*_*w*_ of all “*p*-*p*” gene pairs and, similarly, *μ*_*p*−*n*_, 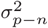 for all “*p*-*n*” gene pairs. Let also *N*_*p*−*p*_ and *N*_*p*−*n*_ denote the number of gene pairs in each category. We use Welch’s two-sample *t*-statistic with unequal variances to quantify the notion of separability:

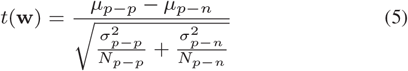

Note that *μ* and *σ*^2^ are functions of **w**, but this dependence is not shown explicitly in equation 5 to keep the notation simple. Maximizing *t*(**w**) is equivalent to minimizing –*t*(**w**). In order to enable sample selection, we also added an L1 regularization term that forces the weights of uninformative samples to zero (Tibshirani, 1996). Our optimization problem then becomes:

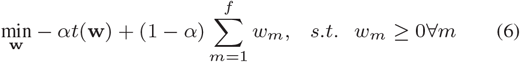

Parameter *α* controls the trade-off between the actual cost and the regularization. The minimization of Equation 6 is done with the Broyden-Fletcher-Goldfarb-Shanno method (Byrd *et al*., 1995).

#### 2.4.1 Global *MLC*

To investigate the effect of creating GO-term specific predictors, we also implemented a version of *MLC* that is applicable to all terms simultaneously. To this purpose, we redefined “*p*-*p*” gene pairs as pairs of two genes which share at least one GO term and “*p*-*n*” pairs as pairs of two genes that share no GO annotations. All the ensuing steps remain the same as for the term-specific *MLC*. We call this method “Global *MLC*” (*MLC*_*G*_).

### 2.5 Experimental set-up

#### 2.5.1 Competing methods

We compared *MLC* to the unweighted *PCC* baseline. To investigate the effect of the use of a term-specific classifier, we created term-specific classifiers from the *PCC* by tuning the classifier parameter *k* individually per GO term and not globally over all terms. We called this approach *PCC*(*k*). We also compared to *GAAWGEFA* which, like *MLC*, learns a weighted co-expression measure (Ray and Misra, 2019). *GAAWGEFA* is not GO-term-specific and optimizes the mean protein-centric precision using a genetic algorithm, so we also constructed a non-term-specific version of *MLC* (*MLC*_*G*_) to compare against. Another way to measure co-expression is the Mutual Rank (*MR*) (Obayashi *et al*., 2018) which is used in the ATTED-II database. Although *MR* neither selects samples nor weighs samples differently, it has been shown to outperform the *PCC* for function prediction (Obayashi *et al*., 2018), so we included it in the comparison as a stronger baseline. Input to *MR* are typically the *PCC* values of all gene pairs, although it can be applied to any co-expression measure. More details on the definition and implementation of each of these methods are given in Supplementary Material 1.

#### 2.5.2 Cross-Validation Experiment

We used the *k*-Nearest Neighbors (*k*-NN) classifier to compare the function prediction performance of the different studied co-expression measures on all *A. thaliana* genes with at least one BP annotation. To counter the imbalance in the dataset, we restricted ourselves to GO terms that annotate at least 1% of the genes. Also, for the weight optimization stage of *MLC*, we randomly sampled an equal number of genes with and without each term. The optimal number of nearest neighbors (*k*) is a parameter of all methods. MLC has an extra regularization parameter *α* (equation 6). We tuned the parameters of MLC independently for each GO term, while we selected the value of *k* that maximized the mean performance over all tested GO terms for the other methods. Parameter tuning was done in a double 3-fold cross-validation loop (Varma and Simon, 2006) using the term-centric *ROCAUC* as performance criterion. The inner loop was used to select the optimal parameter values and the outer loop to evaluate the performance of the tuned models on previously unseen genes (reported as “CV results”). As in this work we are dealing with the problem of identifying which genes should be annotated with a specific GO term, we focus on term-centric evaluation using the mean *ROCAUC*. However, we also compared the methods with two protein-centric measures, the maximum F-measure (*F*_*max*_) and the minimum Semantic Distance (*S*_*min*_) (Supplemental Material 2).

#### 2.5.3 CAFA Experiment

We also evaluated the same methods on the preliminary test set from CAFA3, released by the organizers in June 2017. This dataset contains 6077 training and 137 test genes from *A. thaliana*. After ID mapping, we restricted ourselves only to the genes for which we had both expression data from ArrayExpress and *MR* values from the ATTED database and passed the filtering step described in section 2.1. This left us with 4,889 training genes and 90 test genes, annotated with 707 GO terms. We used the training set to tune the parameters of the tested methods using a 3-fold cross-validation loop. We removed the 10 rarest terms, as there were not enough training and testing genes in all folds. Then, we re-trained each method on the whole training set, using the optimal parameter values found with cross-validation and made predictions for the 697 remaining terms on the 90 test genes (reported as CAFA results). To assess the variability of the results, we performed 1,000 bootstraps, choosing at random with replacement 90 genes at each iteration and re-evaluating the mean term-centric performance. We used these bootstraps to construct 95% confidence intervals.

## 3 Results

### 3.1 All methods outperform the *PCC*

We compared our metric learning approach (*MLC*), as well as Mutual Rank (*MR*) and *GAAWGEFA* to the standard, unweighted PCC using 3fold cross-validation. *PCC* achieved a mean term-centric *ROCAUC* of 0.69, while the performance of both *MR* and *MLC* with the weighted inner product was 0.72 (Table 1). The performance of *GAAWGEFA* was 0.71. *MLC*_*G*_, the non-term-specific version of *MLC*, also achieved a mean *ROCAUC* of 0.72. Although all methods perform fairly similarly according to protein-centric measures (Tables S1-2, Supplementary Material 3), *PCC* performs significantly worse than the other methods on term-centric *ROCAUC* (False Discovery Rate (FDR) < 0.036, Tables S3-5, Supplementary Material 4, effect size 4%). This shows that the *PCC* is indeed a suboptimal co-expression measure.

**Table 1.**
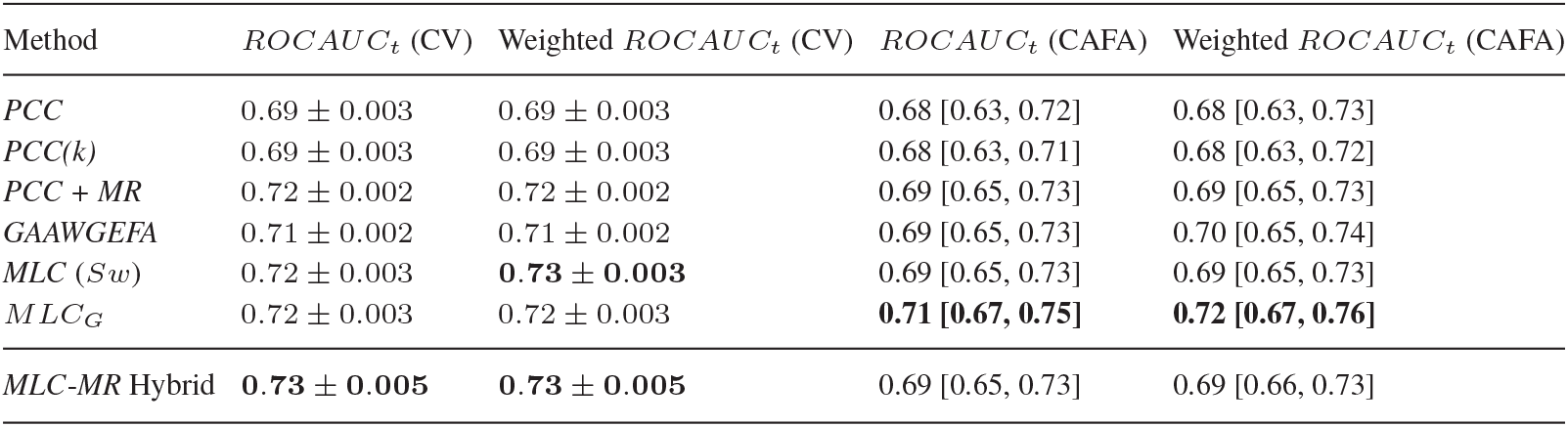
Mean term-centric *ROCAUC* (*ROCAUC*_*t*_) achieved by the methods under comparison using 3-fold crossvalidation (CV, 2nd and 3rd column) and when testing on the CAFA dataset (CAFA, 4th and 5th column). For the cross-validation, we report the average performance over the three folds as well as the corresponding standard error. For the CAFA results we report the performance on the test set as well as the 95% Confidence Intervals from doing 1,000 bootstrapped tests.

### 3.2 *MLC* is the best at predicting specific GO terms

Although *MR*, *GAAWGEFA* and *MLC* perform equally on average, one is typically not interested in predicting GO terms that are “near” the ontology root, as most of them describe too general biological processes (Clark and Radivojac, 2013). Therefore, we compared the performances of these methods as a function of term specificity. As measures of specificity, we used the maximum path length to the ontology root and the Resnik Information Content (IC) (Resnik, 1995). One way to take term specificity into account is to calculate the weighted term-centric *ROCAUC*, where each term is weighted by its IC when calculating the average. As shown in Table 1, *MLC* achieves the highest weighted *ROCAUC*. The difference is statistically significant for all methods except for *MR* (Table S6, Supplementary Material 4), although the effect size is small (1.5%).

Furthermore, we grouped the GO terms into quintiles (quantiles at 0, 20,40,60 and 80%) and plotted the distribution of the percent differences in performance of *MLC* from *MR* for each quintile (Figure 2a). We observed that for the first two quintiles (i.e. the 40% most frequent terms), *MLC* performs worse than *MR*, while for the 60% most specific terms, both the mean and the median performance of *MLC* is better (Figure 2a). Further analysis showed that for the very general terms, *MLC* makes a lot more type I errors (false positives) than for the more specific ones (Figure S1, Supplementary Material 5) and that makes it underperform with respect to *MR*. The Spearman correlation between percent difference and Resnik IC was 0.26. The same pattern is evident when comparing *MLC* to all other methods (*PCC*, *GAAWGEFA* and *MLC*_*G*_), as well as when replacing Resnik IC with the path length to the ontology root (Tables S7-8, Figures S2-3, Supplementary Material 6). From that we can conclude that term-specific *MLC* is the preferred method for finding genes belonging to rarer terms.

**Figure 2.**
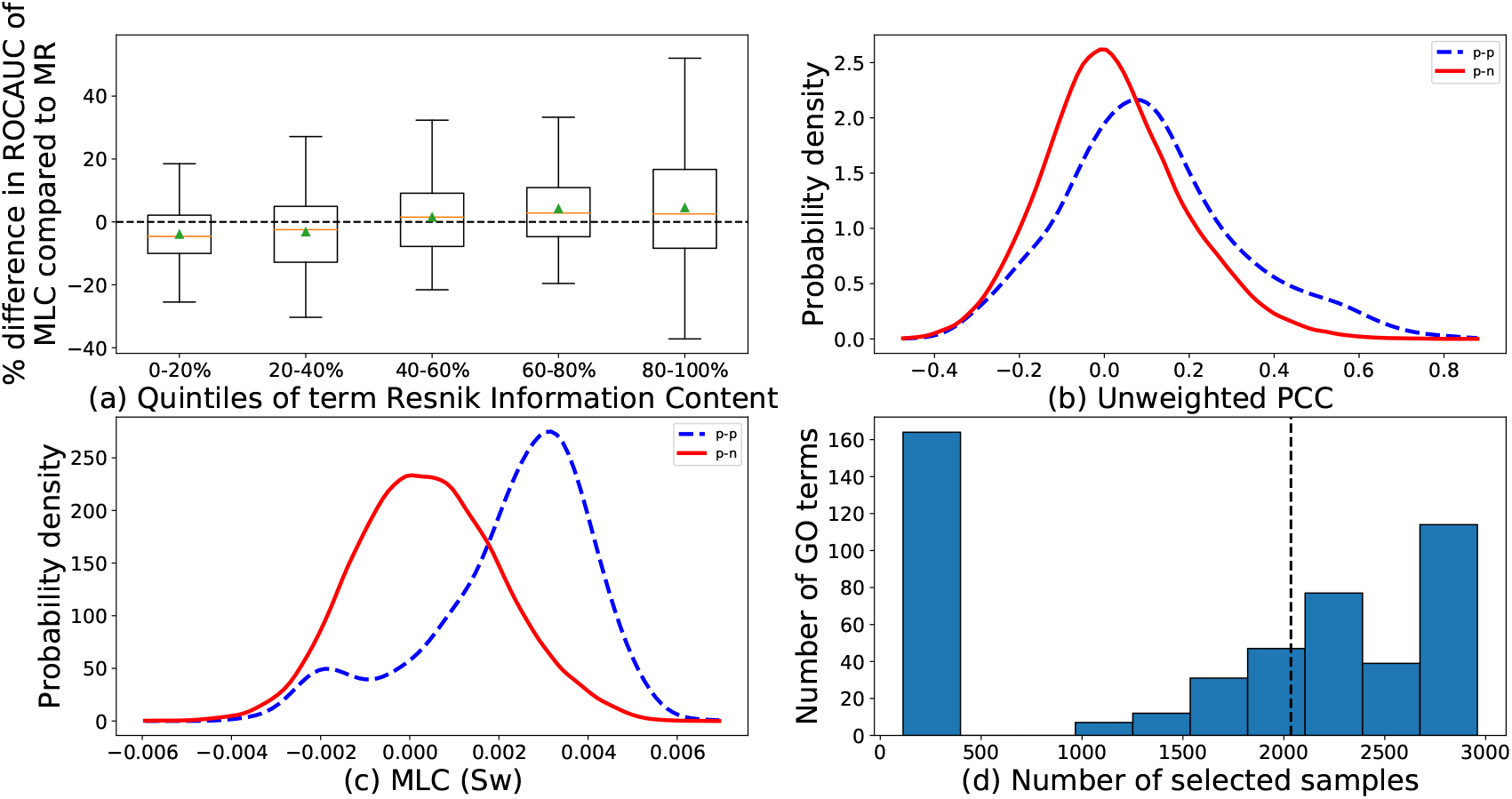
(a)Percent increase in *ROCAUC* of *MLC* (*S*_*w*_) with respect to *MR* as a function of Resnik Information Content. For each set of terms in each quintile of Information Content, the corresponding box includes the two middle quartiles of the percent increase for these terms. An orange line denotes the median. The error bars extend to 1.5 times the range of the two middle quartiles and outlier points are shown as dots. (b-c) Distributions of co-expressions for genes annotated with term GO:1903047. In dashed blue lines, the co-expression values between a test and a training gene that both are annotated with that term. In solid red lines, the co-expression between test genes annotated with that GO term and training genes that are not. Co-expression is measured as the PCC (b) and the *S*_*w*_ trained by MLC (c). The *x*-axis shows the co-expression values and the *y*-axis the probability density estimated with Gaussian kernels. Note that the PCC and *S*_*w*_ have different ranges due to the weight optimization. (d) Histogram of the number of samples that were selected for each GO term. The *x*-axis corresponds to the number of selected samples and the *y*-axis to how many GO-term-specific similarity functions selected that many samples. The dashed line denotes the median number of non-zero weights.

### 3.3 *MLC* tunes the weights to find “*p*-*p*” pairs

The goal of *MLC* is to choose the weights so that for test genes that have a particular GO annotation, the learned similarities are higher to training genes that have the same annotation than to genes that do not. As an example, Figures 2a and b show the distribution of co-expression similarities between the test genes annotated with term GO:1903047 (mitotic cell cycle process) and all training genes for the *PCC* and *MLC* similarities respectively. It is clear that for *MLC*, the test genes are a lot more similar to training genes with the same annotation. Note, however, that, for this term, a significant portion of the similarities are negative (small blue peak in Figure 2b). This means that some positive genes are anti-correlated to the rest. For these cases *MLC* will make Type II errors (false negatives).

Figure 2d shows the distribution of the number of selected samples for each GO term. For about 33% of all GO terms, *MLC* selected less than 9% of the available samples (252 or less), setting all other weights to zero. The median number of selected samples was 2,035 out of 2,959 or about 69% of all samples. Moreover, for about 23% of the terms, MLC kept all the samples and weighted them more or less equally (maximum standard deviation of weights = 0.006), in which case MLC was almost equivalent to the unweighted inner product. However, MLC still had significantly better performance for those terms than baseline *PCC* (median difference 0.03, p-value =0.002, Wilcoxon rank sum test). We also found that individually tuning *k* per GO term for the PCC gave on average the same term-centric *ROCAUC* as the baseline *PCC* (*PCC* (*k*), Table 1), so the performance improvement is not caused by simply choosing the optimal *k* value for each GO term. Finally, we observed a small negative correlation between term Information Content and the number of samples selected (Spearman *ρ* = –0.09, p-value = 0.057). This means that *MLC* has a slight tendency to select fewer samples for more specific terms, but this result is not statistically significant.

### 3.4 The weights learned by MLC help at identifying relevant experimental conditions

The weights learned by *GAAWGEFA* are roughly uniformly distributed between 0 and 1 (Kolmogorov-Smirnov test statistic = 0.011, p-value = 0.847) and are not correlated to any of the term-specific weight profiles of *MLC*, which tend to have an exponential-like distribution (Figures S4-5, Supplemental Material 7), as many samples get a weight of zero. Continuing on the example from before, for term GO:1903047 (mitotic cell cycle process) *MLC* gives the highest weight to a sample of a plant grown in the absense of phosphorus, which has been shown to restrict the cell division rate (Kavanová *et al*., 2006). Among the samples with highest weights are also many samples from experiments studying seed germination, a process closely linked to cell cycle (Vázquez-Ramos and de la Paz Sánchez, 2003). Finally, two *IBM1* mutant samples were selected with very high weights. The *IBM1* gene codes for histone demethylation protein and has GO annotations that include flower, root and pollen development. The complete weight profile is shown in Figure S6 (Supplemental Material 8). This example shows that *MLC* is able to identify RNA-Seq samples that are relevant to the process under study and that by examining the weight profile of *MLC*, one can interpret its predictions more easily.

### 3.5 The weights learned by MLC are consistent with the ontology structure and the existing annotations

We compared the sample weights learned by MLC between parent-child GO term pairs in the following three cases: 1) the parent has only one child term and both parent and child annotate exactly the same genes, 2) the parent has only one child term but annotates more genes than its child, and 3) the parent has exactly two children, meaning that the parent annotates the union of the genes of its children. In the first case, the weight profiles for parent and child are identical (mean Pearson correlation of 1). In the second case, the profiles are similar but not identical (mean Pearson correlation of profiles 0.47). The difference in mean similarity between the two groups is statistically significant (permutation p-value < 10^−5^). Furthermore, the larger the difference in number of extra genes of the parent term, the smaller the profile correlation (Spearman *ρ* = –0.53, *CI*_95%_ = [–0.65, –0.40]). The profile similarities are even smaller in the third case (mean of 0.36) and significantly smaller than those of case 2 (permutation p–value =0.0002). This is expected as in this case the parent contains two distinct sets of genes that correspond to two different biological processes. Again, we found a negative correlation between the number of different genes and the profile similarity of pairs (Spearman *ρ* = –0.47, *CI*_95%_ = [–0.61, –0.31]).

To generalize this finding, we hierarchically clustered the GO terms (complete linkage, Jaccard distance between the gene sets associated with each GO term cutoff of 0.6). The resulting clusters are shown in Figure 3 along with the pairwise distances of the GO terms. 64 out of 176 clusters contained at least three terms. For each of these 64 clusters, we randomly sampled 10,000 equal-sized sets of GO terms and calculated the mean pairwise similarity in those sets to calculate a permutation p-value. For 62 out of these 64 clusters we found that the pairwise weight-profile similarities of their members (Figure S2, SM2) are significantly higher than random with a False Discovery Rate of 0.05. Also, pairwise profile similarities are positively correlated with the pairwise Resnik semantic similarity of GO terms (Spearman *ρ* = 0.16, *CI*_95%_ = [0.15, 0.17]). Based on these observations, we conclude that sample weights reflect the gene annotations of each term.

**Figure 3.**
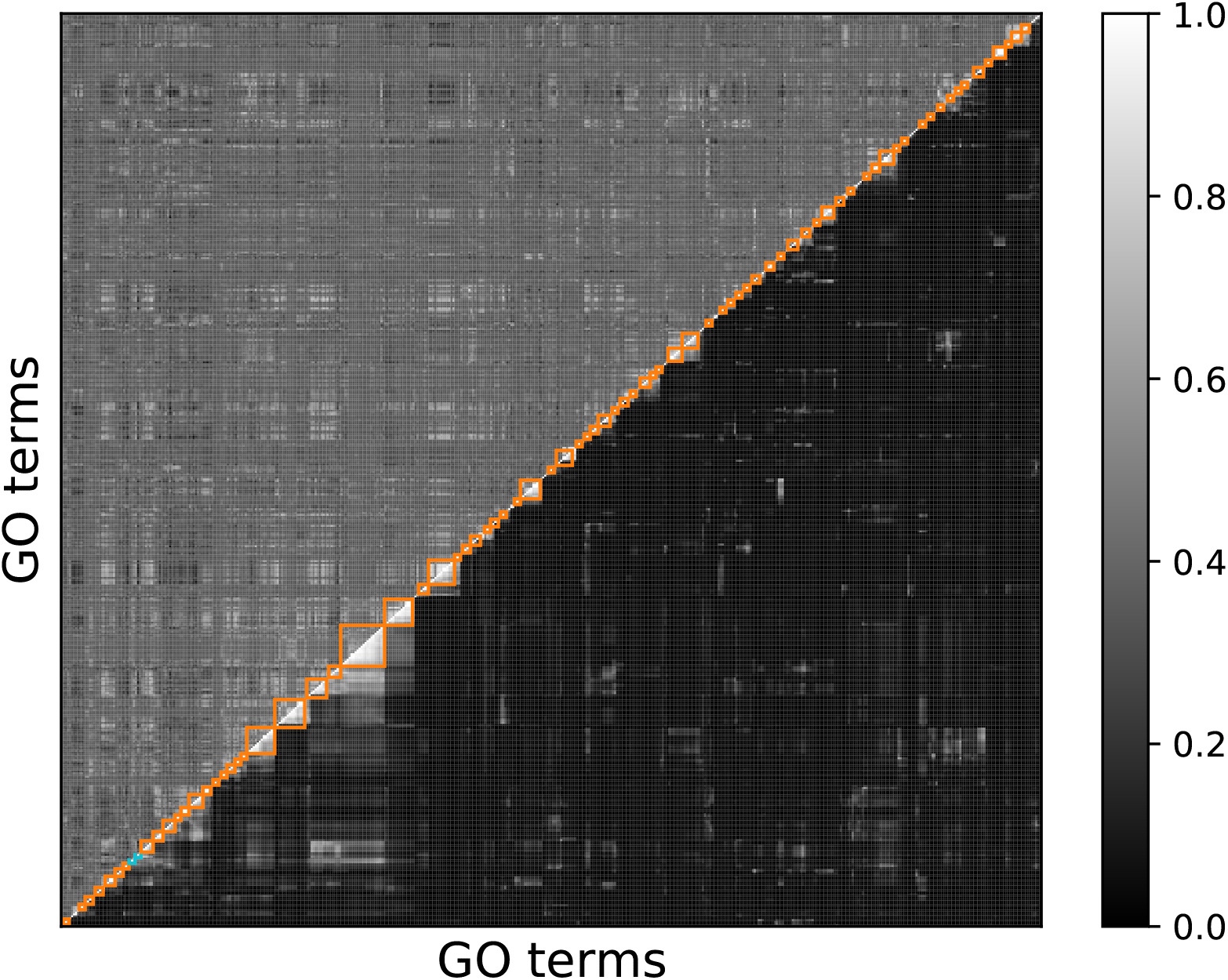
Pairwise Jaccard similarities (below the anti-diagonal) and weight profile similarities (Pearson correlations, above the anti-diagonal) of the tested GO terms. Both the *x* and *y* axes show the GO terms ordered so that terms in the same cluster are adjacent. The grey-scale indicates the similarity, with dark being low and bright being high similarity. The profile correlations have been scaled so that they are in the range [0, 1]. The squares highlight the clusters containing at least 3 terms, (cutting the dendrogram at a Jaccard distance threshold of 0.6 when using complete linkage). Light blue boxes indicate the clusters that are not significantly enriched with terms with similar weights and orange-colored clusters are significantly enriched after FDR correction.

### 3.6 Using all samples obscures co-expression

Next, we investigated the terms for which *MLC* performed sample selection, i.e. assigning a non-zero weight to at most 9% of the samples. We looked at the *PCC* values for “*p*-*p*” and “*p*-*n*” gene pairs. Figure 4a shows an example of the distributions of the *PCC* values for “*p*-*p*” and “*p*-*n*” pairs for term “GO:1903047” (mitotic cell cycle process). Next, we calculated the *PCC* for all “*p*-*p*” and “*p*-*n*” pairs, but only considering the samples that were selected by *MLC* (i.e. had a non-zero weight assigned), as well as only considering the samples that were not selected by *MLC* (i.e. were assigned a weight of zero), shown in figures 4b-c respectively. One can notice that the two distributions (“*p*-*p*” and “*p*-*n*” *PCC* values) differ more when taking the *MLC* selected samples in consideration (compare Figure 4a with Figure 4b). When only considering the samples that were not selected, the two distributions differ in a similar way to when all samples are being considered (compare Figure 4a with Figure 4c).

**Figure 4.**
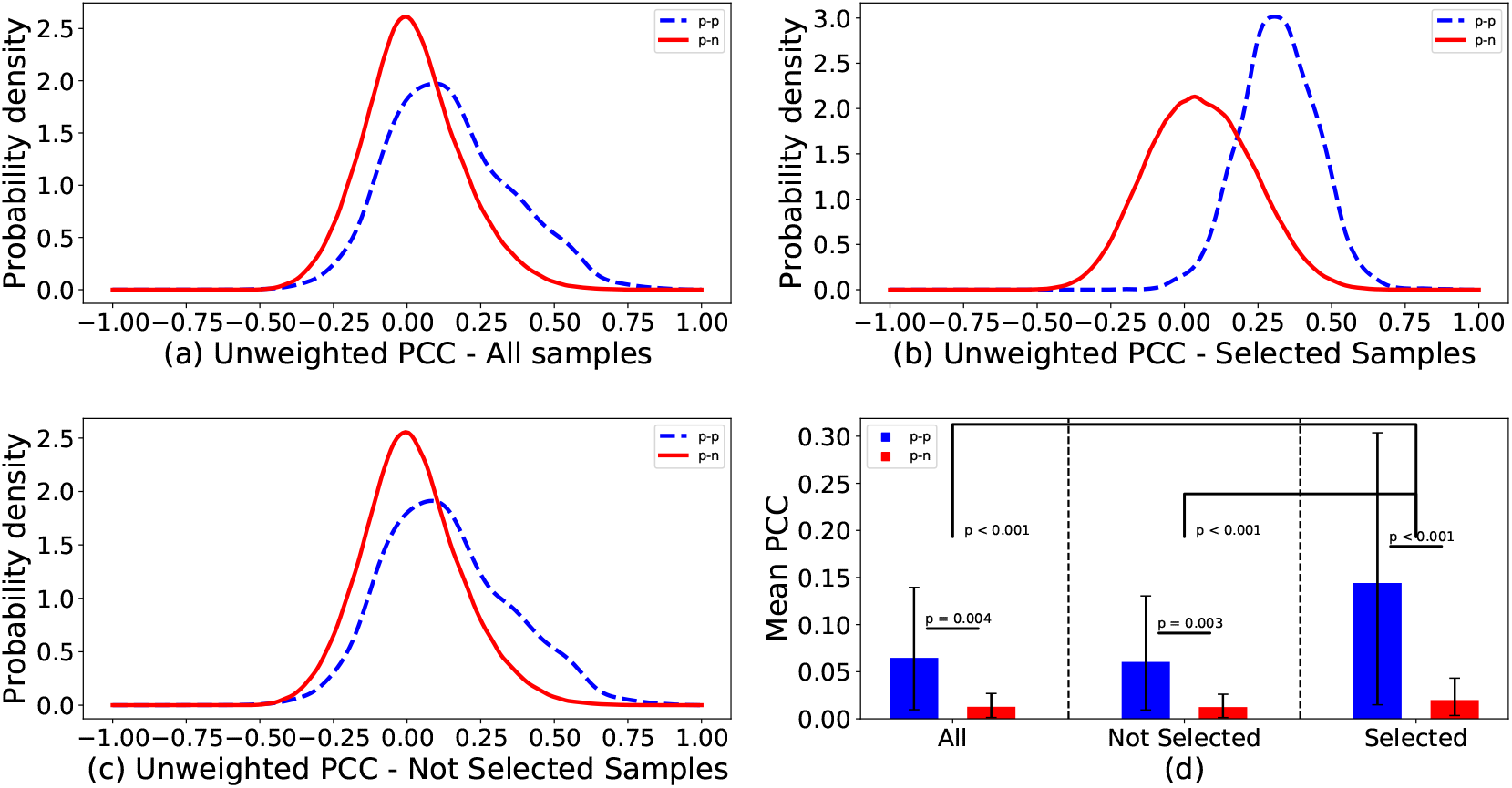
(a-c): Distributions of Pearson correlations for pairs of training genes that are both annotated with term GO:1903047 (“*p*-*p*”, blue dashed), and for pairs of training genes of which only one is annotated with that term (“*p*-*n*”, red solid). The correlations are calculated using all samples (a), the samples that were selected by MLC (b) and the samples that were not selected (c). (d): The means of the distributions of (a-c), over all GO terms where less than 10% of the samples where used to calculate the co-expression (more than 90% zero weights). Values for “*p*-*p*” pairs are colored blue and for “*p*-*n*” pairs red. The error bars show the 95% confidence intervals for the means, calculated with 1,000 bootstraps. Above the bars the bootstrap p-values are shown for the pairwise comparisons.

For every GO term for which *MLC* performs sample selection, we calculated the mean *PCC* of all “*p*-*p*” and “*p*-*n*” pairs under the three sample sets (all samples, not selected samples and selected samples). Figure 4d shows the average of these values of all GO terms. We also performed 1,000 bootstraps, sampling GO terms with replacement to obtain 95% confidence intervals for these averages. We observed that the difference in mean co-expression between “*p*-*p*” and “*p*-*n*” in the samples not selected by *MLC* is similar to the difference in all the samples. Although these differences are statistically significant, they are also significantly smaller than the difference in the *MLC*-selected samples (bootstrap p-value <0.001) (Figure 4d). This means that although the whole dataset does contain a few samples that are informative for these GO terms, calculating the co-expression over a larger set of samples can corrupt the “real” co-expression signal, increasing the difficulty of discovering new genes that play a role in these processes.

### 3.7 Combining Mutual Rank and *MLC*

The performances of *MLC* and *MR* are positively correlated (Spearman *ρ* = 0.13, p-value =0.003). We also applied *MR* on the co-expression similarities obtained with *MLC*, as *MR* is in principle not restricted to using only the *PCC*. We found a small improvement compared to standalone *MLC*, with a mean ROC AUC of 0.73. Also, the performances of *MLC* and *MLC* + *MR* were highly correlated (Spearman *ρ* = 0.97, p-value ≪ 10^−20^). We tried another approach to combine *MLC* and *MR* depending on the performance of the methods. If the training *ROCAUC*_*t*_ of *MR* was larger than 0.8 for a GO term, we used the predictions of *MR* for that term, otherwise we used the predictions of *MLC*. This combined classifier had an incrementally larger term-centric *ROCAUC* (0.73, Table 1 - Hybrid), though statistically significant (p-value = 0.008, two-sample t-test). The threshold of 0.8 training *ROCAUC* was chosen arbitrarily and was not tuned to maximize performance. This naïve hybrid classifier shows that there is potential to improve performance by combining *MLC* and *MR* in more sophisticated ways.

### 3.8 CAFA Results

Lastly, we benchmarked MLC on 90 temporary *A. thaliana* targets from the CAFA3 competition. The results are similar, but the small size of the dataset does not allow us to draw any meaningful conclusions (Table 1). Both MR and MLC outperform PCC on average, but the confidence intervals are much wider.

## 4 Discussion

### 4.1 MLC

We introduced *MLC*, a metric learning method for building automatic function predictors from a large collection of expression data. *MLC* calculates gene co-expression by assigning GO-term-specific weights to each sample. The weights aim at maximizing the co-expression similarity between genes that are annotated with that GO term. In general, training GO-term specific classifiers has the disadvantage that individual classifiers fail to see the “bigger picture” and cannot exploit the correlations between terms imposed by the ontological structure. However, we showed that the weight profiles learned by *MLC* do correlate with real biological knowledge such as semantic similarity in the ontology graph and gene annotation similarity. Due to the use of the L1 regularization, *MLC* can also select informative samples by setting the weights of non-informative samples to zero, but even for the terms where no selection is performed, *MLC* can weigh the samples differently leading to an improvement in performance compared to *PCC*. Moreover, we showed that the samples that are selected come from biological conditions relevant to the GO term in question.

Our method is designed to work well with a Guilt-By-Association approach like the *k*-NN classifier. This classifier assigns a GO term to a test gene if a large enough fraction of its top co-expressed training genes are annotated with that term. To achieve this, *MLC* tries to maximize the difference between the average co-expression between gene pairs that are both annotated with the GO term of interest (“*p*-*p*” pairs) and the average co-expression between gene pairs only one of which is annotated with the term (“*p*-*n*” pairs). During the training phase, our model ignores gene pairs where neither gene has the term of interest (“*n*-*n*” pairs). Such pairs could either include two genes that have common GO annotations, but different from the GO term of interest or two genes with completely different annotations. For the first case, one might be tempted to think that the co-expression of such pairs should be high. However, if their common function is different from the term of interest, it is likely that they are correlated for another set of samples than the one related to the GO term of interest, and, consequently, are thus uninformative for that GO term. For the second type of “*n*-*n*” pairs, the ones that share no annotations whatsoever, it might make sense to want their co-expression to be 0, as they are expected to be dissimilar over any set of samples. However, we decided to ignore these pairs as they do not add any term-specific information, so it is not clear how they will affect the identification of samples specifically relevant for a specific term. This might be problematic as for a negative test gene (i.e. a gene that should not be annotated with the GO term of interest) we cannot exclude that it can be as highly co-expressed to positive as to negative genes, because we did not tune the co-expression values for “*n*-*n*” pairs. For very frequent terms with a lot of positive training genes this leads to a lot of false positive predictions, which might explain the poor performance of *MLC* for frequent terms.

The similarity function that we used as a basis for *MLC* is the weighted inner product (*S*_*w*_). We chose this measure because its unweighted version is identical to the unweighted PCC for centered and scaled data, but it has a simpler form which eases the computational burden. The weighted versions of the inner product and PCC are no longer identical, as the data are no longer scaled after weighing the samples. This has as side-effect that the similarity functions that MLC learns are not necessarily in the range [−1,1], like the PCC. In most cases, their range is much narrower as can be seen in Figure 2b for GO:1903047. Also, because of the range differences, it is not trivial to compare the similarity of two genes across different GO terms. For the purpose of classification with the *k*-Nearest Neighbors classifier, however, the range of the metric is insignificant (only the relevant rankings are important to find the proper neighborhood).

Our model is more general and not restricted to only the inner product, though. The main idea is to maximize the difference between the similarity of *p*-*p* and *p*-*n* pairs. This is done by maximizing the *t*-statistic between the two distributions of similarities. This means that MLC can also be applied to any measure of similarity such as the weighted PCC, weighted Spearman correlation, Euclidean distance etc.. Regardless of the chosen metric, the two classes (“*p*-*p*” and “*p*-*n*”) do not meet the assumptions for applying Student’s *t*-test, as the similarity values are neither normally distributed nor independent. This is not an issue, though, because we do not use the t-statistic to compute a p-value (exploit that the t-statistic is distributed according the Student’s t-distribution under these assumptions), but only to quantify the class separability (Theodoridis and Koutroumbas, 2008). Equivalently, we could have used any other measure of class separability, for instance the Fisher Discriminant Ratio (Fisher, 1936) or the Davies-Bouldin index (Davies and Bouldin, 1979).

### 4.2 Comparison to related methods

Our work validates the observation that *PCC* is not the optimal co-expression measure for AFP. The Mutual Rank (*MR*) attempts to obtain more robust and noise-free co-expression values by converting the *PCC* values into ranks and averaging the reciprocal rankings of two genes (Obayashi *et al*., 2018). *MLC* takes a fundamentally different approach, operating on the sample level rather than the correlation level. First and foremost, as we mentioned above, it removes samples that do not help at discriminating between genes that do or do not perform a certain function. With that MLC gives insight into which samples are important for a given GO term, which subsequently can be used to investigate the expression patterns of the GO term related genes across these samples. Weighing samples differently can also be viewed as a way of denoising. For example it can compensate for the issue that an expression change of 1 unit has a different meaning in different samples due to technical variations, such as for example differences in sequencing depth or sample preparations. Our results have shown that *MLC* is more beneficial than the *MR* approach for the more specific - and arguably more useful - GO terms.

A similar method to *MLC* is *GAAWGEFA*, which learns a weight for each sample in a dataset adnd then applies a weighted Pearson correlation. There are two fundamental differences between the two methods. Firstly, *GAAWGEFA* aims at good protein-centric performance, i.e. it tries to do well on average for all genes and therefore learns only one set of sample weights. On the other hand, *MLC* aims at maximizing the performance for each GO term individually. Secondly, *GAAWGEFA* learns the weights using a genetic algorithm. For *MLC*, we used the inner product, which allowed us to have a simple optimization problem that can be solved very efficiently. Even though *MLC* has to be run for each term separately, it is still 67% faster than *GAAWGEFA* and, unlike *GAAWGEFA*, runs for different GO terms can be carried out in parallel to achieve even greater speed-up. Next to those differences, *MLC* makes more accurate predictions for rarer terms and provides interpretability of the predictions by examining the term-specific sample weight distributions. Furthermore, in the context of selecting expression samples a related technique is biclustering. Biclustering is an umbrella term for a diverse set of algorithms that simultaneously select subsets of genes and samples, so that the genes in the same subset (bicluster) have similar expression to each other within the samples of that bicluster. It is typically expected that each bicluster reflects a biological process and that makes the rationale of *MLC* appear similar to a biclustering approach. Although both approaches make use of sample selection and aim at discovering genes involved in the same biological processes, they are fundamentally different in the sense that *MLC* is supervised and biclustering unsupervised. Biclustering does not make use of GO annotations, but only of the expression matrix. Often, observing enrichment of certain GO terms or KEGG pathways in the genes of biclus-ters is one of the ways to validate a biclustering result (Santamaría *et al*., 2007). On the other hand, *MLC* starts with a set of genes whose GO annotations are known (or at least partly known) and tries to use the expression matrix in order to identify which of the remaining genes participate in a particular biological process by defining a co-expression measure specific to that process.

### 4.3 Possible Extensions

*MLC* learns the sample weights automatically from the available data and does not rely on information about the samples’ biological condition or tissue. As curation efforts increase and the amount of well-annotated data in public databases grows larger with time, in the future it might be useful to extend *MLC* to incorporate such knowledge. A possible way to do that would be a group LASSO approach (Yuan and Lin, 2006). Group LASSO uses predefined groups of samples and forces the weights of all samples in a group to be equal. Each such group could contain technical and biological replicates, samples from the same tissue or samples from similar knockout experiments and perturbations.

*MLC* does not account for the possibility that genes that show exactly opposite expression patterns (i.e. genes with large negative correlation) might also be involved in the same biological process. Future work should show whether this is useful for AFP. Finally, in this work, we applied *MLC* on finding candidate genes for GO terms from the BPO. However, it can be useful for any gene annotation problem that can be solved with expression data, such as finding members of KEGG pathways or genes that are likely to influence a given phenotypic trait. As *MLC* is computationally efficient it can easily be applied to a large number of different terms/phenotypes, offering state-of-the-art performance with the added benefit of allowing users to understand which parts of the dataset influence the predictions.

