## Supplementary material for "Metric Learning on Expression Data for Gene Function Prediction": Supplemetary Material

Stavros Makrodimitris [1,2,*], Marcel J.T. Reinders [2,3] and Roeland C.H.J. van Ham [1,2]

[1]Delft Bioinformatics Lab, Delft University of Technology, Delft, the Netherlands,
[2]Keygene N.V., Wageningen, the Netherlands and
[3]Leiden Computational Biology Center, Leiden University Medical Center, Leiden, the Netherlands.
[*]To whom correspondence should be addressed.

**3 Protein-Centric Results** **4**

**4 Statistical Significance of Differences in Cross-Validation Performance** **5**

**5 False positive rates for general and specific terms** **6**

**6 Performance as a function of term specificity** **7**

**7 Comparison of *GAAWGEFA* and *MLC* weights** **12**

**8 Weight profile** **13**

### 1 Competing Methods

#### 1.1 Mutual Rank (*MR*)

An alternative way to measure co-expression is using the Mutual Rank (*MR*) which has been successfully used in the ATTED-II co-expression database [1]. Although *MR* neither selects samples nor weighs samples differently, it has been shown to outperform the *PCC* for function prediction. Input to *MR* are typically the *PCC* values of all gene pairs, although it can be applied to any co-expression measure. For every gene i, the co-expression values to all other genes are ranked in descending order. We use $rank_i(j)$ to denote the rank of gene $j$ in terms of similarity to $i$. Note that $rank_i(i) = 0$. The *MR* value between two genes is calculated as the geometric mean of $rank_i(j)$ and $rank_j(i)$:

$$MR(\mathbf{x}_i, \mathbf{x}_j) = \sqrt{rank_i(j) \times rank_j(i)} \quad (1)$$

*MR* is expected to be more robust to spurious correlations caused by outliers (due to the geometric mean) and can also handle cases where a gene tends to have very high or very low correlations with many other genes, which bias the co-expression ranks leading to poorer performance for the $k$-NN classifier. We downloaded the pairwise *MR* values constructed from public RNA-Seq data for all *A. thaliana* genes from the ATTED-II website (`http://atted.jp/top_download.shtml`).

#### 1.2 *GAAWGEFA*

For *GAAWGEFA* [2] we used the MATLAB code provided by the authors at `http://sampa.droppages.com/GAAWGEFA.html`. Because of the long runtime we did not tune any of the parameters of the algorithm and used the default settings as mentioned in the paper:

- Initial population size: 20
- Crossover Probability: 0.85
- Mutation Probability: 0.1
- Maximum iterations: 1000

After the 150-th iteration we enabled the extra option of early termination, i.e. we stopped the optimization if the fitness function had not increased by at least 1% between two consecutive generations. We did tune the number of nearest neighbors used in the $k$-NN classifier as previously.

### 2 Evaluation Measures

#### 2.1 Unweighted and Weighted term-centric ROC AUC

In the setting used in this work, we are interested in finding the genes that perform a given function, so term-centric evaluation is the most interesting. As in the CAFA papers, we use the term-centric area under the ROC curve ($ROCAUC$). For every GO term $l$, we compute a $ROCAUC$ score for the binary problem of finding all test genes that are annotated with that term and then we report the average of that value over all GO terms. We also use the weighted version of this measure ($ROCAUC_w$), in which we calculate a weighted average, weighing each term by its Resnik Information Content ($IC(l)$) [3].

#### 2.2 Protein-centric Measures

For protein-centric evaluation, we use the F1 score ($F_{max}$) and the Semantic Distance ($S_{min}$) [4]. Similar to the CAFA evaluation, we start with the posterior probabilities for each pair of test gene and GO term. As these measures require hard predictions (i.e. either 0 or 1, not a posterior probability) we compute them for 21 equally-spaced thresholds from 0 to 1 (i.e. in steps of 0.05). We then only report the metric value at the optimal threshold for each measure (the maximum for the F1 and the minimum for the Semantic Distance) and for each evaluated algorithm [5].

### 3 Protein-Centric Results

We compared 5 methods: *PCC*, *PCC+MR*, *GAAWGEFA*, *MLC* and $MLC_G$ using 2 term-centric and 2 protein-centric evaluation measures. Term-centric performance is shown in Table 1 in the main document. Tables S1 and S2 show the protein-centric $F_{max}$ and $S_{min}$ for these methods on the cross-validation and CAFA3 datasets respectively. We observe that, according to both metrics, all methods achieve more or less equivalent protein-centric performance, with the exception of the *PCC*, which performs significantly worse (section 4). The *MR* seems to be consistently the top method, but the differences are very small and not statistically significant (See also section 4).

Table S1: Comparison of the protein-centric $F_{max}$ and $S_{min}$ of the tested methods using 3-fold cross-validation. For each method and metric the mean and standard error are shown. Higher $F_{max}$ and **lower** $S_{min}$ denote better performance.

| Method | $F_{max}$ | $S_{min}$ |
|---|---|---|
| *PCC* | $0.34 \pm 0.001$ | $19.12 \pm 0.11$ |
| *PCC + MR* | $0.36 \pm 0.001$ | $18.85 \pm 0.11$ |
| *GAAWGEFA* | $0.35 \pm 0.002$ | $18.97 \pm 0.10$ |
| *MLC* ($S_w$) | $0.35 \pm 0.001$ | $18.89 \pm 0.13$ |
| $MLC_G$ ($S_w$) | $0.35 \pm 0.003$ | $19.00 \pm 0.17$ |

Table S2: Comparison of the protein-centric $F_{max}$ and $S_{min}$ of the tested methods using the CAFA3 data. For each method and metric the mean and 95% confidence interval are shown. Higher $F_{max}$ and **lower** $S_{min}$ denote better performance.

| Method | $F_{max}$ | $S_{min}$ |
|---|---|---|
| *PCC* | 0.25 [0.22, 0.29] | 21.27 [18.78, 29.17] |
| *PCC + MR* | 0.27 [0.24, 0.30] | 21.18 [18.78, 28.99] |
| *GAAWGEFA* | 0.25 [0.22, 0.29] | 21.32 [18.90, 29.10] |
| *MLC* ($S_w$) | 0.26 [0.23, 0.29] | 21.56 [18.87, 29.11] |
| $MLC_G$ ($S_w$) | 0.27 [0.24, 0.31] | 21.27 [18.81, 29.22] |

### 4 Statistical Significance of Differences in Cross-Validation Performance

We use four evaluation metrics ($ROCAUC$, $ROCAUC_w$, $F_{max}$, $S_{min}$) to compare 5 methods ($PCC$, $MR$, $GAAWGEFA$, $MLC$ and $MLC_G$). We used the paired-sample $t$-test to compare all 10 different pairs of methods across the three cross-validation folds. This test tests the null hypothesis that the mean difference in performance between two methods across the three folds is not different from zero. For every combination of two methods and a metric we obtained a p-value, so in total we obtained 40 p-values. We corrected all these p-values jointly for multiple testing using the Benjamini-Hochberg method for controlling the False Discovery Rate (FDR). The results are listed in tables S3 to S6. $MR$ and $MLC$ are significantly better than $PCC$ according to all four metrics.

Table S3: FDR-corrected p-values for the null hypothesis that the cross-validation $ROCAUC$ of two methods is not different from zero. Entries are colored in red if the row method is significantly worse than the column method (FDR $<$ 0.05) and in green if the row method is significantly better than the column method (FDR $<$ 0.05).

| | $PCC + MR$ | $GAAWGEFA$ | $MLC$ ($S_w$) | $MLC_G$ ($S_w$) |
|---|---|---|---|---|
| $PCC$ | 0.008 | 0.038 | 0.021 | 0.041 |
| $PCC + MR$ | | 0.088 | 0.272 | 0.180 |
| $GAAWGEFA$ | | | 0.066 | 0.175 |
| $MLC$ ($S_w$) | | | | 0.372 |

Table S4: FDR-corrected p-values for the null hypothesis that the cross-validation $ROCAUC_w$ of two methods is not different from zero. Entries are colored in red if the row method is significantly worse than the column method (FDR $<$ 0.05) and in green if the row method is significantly better than the column method (FDR $<$ 0.05).

| | $PCC + MR$ | $GAAWGEFA$ | $MLC$ ($S_w$) | $MLC_G$ ($S_w$) |
|---|---|---|---|---|
| $PCC$ | 0.008 | 0.038 | 0.016 | 0.041 |
| $PCC + MR$ | | 0.088 | 0.229 | 0.189 |
| $GAAWGEFA$ | | | 0.041 | 0.178 |
| $MLC$ ($S_w$) | | | | 0.155 |

Table S5: FDR-corrected p-values for the null hypothesis that the cross-validation $F_{max}$ of two methods is not different from zero. Entries are colored in red if the row method is significantly worse than the column method (FDR $<$ 0.05) and in green if the row method is significantly better than the column method (FDR $<$ 0.05).

| | $PCC + MR$ | $GAAWGEFA$ | $MLC$ ($S_w$) | $MLC_G$ ($S_w$) |
|---|---|---|---|---|
| $PCC$ | 0.038 | 0.113 | 0.038 | 0.155 |
| $PCC + MR$ | | 0.154 | 0.302 | 0.047 |
| $GAAWGEFA$ | | | 0.258 | 0.180 |
| $MLC$ ($S_w$) | | | | 0.180 |

Table S6: FDR-corrected p-values for the null hypothesis that the cross-validation $S_{min}$ of two methods is not different from zero. Entries are colored in red if the row method is significantly worse than the column method (FDR $<$ 0.05) and in green if the row method is significantly better than the column method (FDR $<$ 0.05).

| | $PCC + MR$ | $GAAWGEFA$ | $MLC$ ($S_w$) | $MLC_G$ ($S_w$) |
|---|---|---|---|---|
| $PCC$ | 0.021 | 0.066 | 0.038 | 0.155 |
| $PCC + MR$ | | 0.119 | 0.223 | 0.138 |
| $GAAWGEFA$ | | | 0.246 | 0.181 |
| $MLC$ ($S_w$) | | | | 0.229 |

#### 5 False positive rates for general and specific terms

To explain why $MLC$ performs better for specific terms than for general ones, we compared the ROC curves of the 20% most specific terms to those of the 20% most general ones. Term specificity was measured by the Resnik Information Content [3]. To get an indication of the general behaviour pattern of $MLC$, we averaged all ROC curves in each of the two groups of GO terms. Figure S1 shows these two average curves. We observed that near the point $(0, 0)$, which corresponds to the genes that $MLC$ classified as positive with high posterior probability, the average ROC curve of the specific terms is increasing a lot more sharply than the one of the general terms which is smoother. This means that genes for which $MLC$ is most confident that are positive are indeed enriched with positive labels for the specific terms. The curve of the general terms is increasing more smoothly, meaning that for those terms, $MLC$ is scoring a lot of negative genes highly, resulting in more false positive predictions.

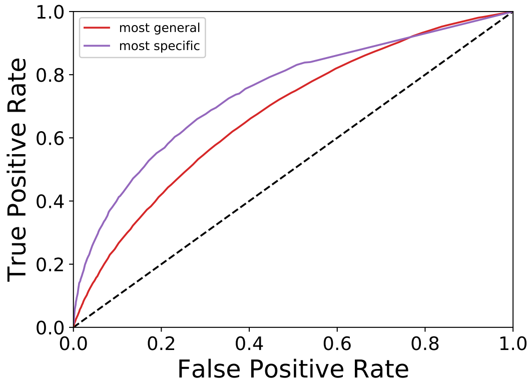

Figure S1: Term-centric ROC curves averaged over the terms with the 20% highest IC (purple) and with the 20% lowest IC (red). The $y = x$ line which represents the expected ROC of a random classifier is shown with a black dashed line.

### 6 Performance as a function of term specificity

For each pair of methods, we calculate the percent difference in $ROCAUC$ between them for each GO term. Tables S7 and S8 show the Spearman correlation of these values with the Information Content and the maximum path length to the ontology root of each term respectively. Moreover, in Figures S2 and S3 we plot these differences for each pair. From these we can clearly conclude that term-specific $MLC$ is the best of all tested methods at predicting specific terms.

Table S7: Spearman correlation of the % difference in performance between the method in each row and the one each column with term Resnik Information Content. Statistically significant values (FDR $<$ 0.05) are shown in bold.

| | $PCC$ | $PCC + MR$ | $GAAWGEFA$ | $MLC$ | $MLC_G$ |
|---|---|---|---|---|---|
| $PCC$ | - | -0.028 | -0.041 | **-0.174** | -0.055 |
| $PCC + MR$ | 0.028 | - | 0.016 | **-0.165** | 0.029 |
| $GAAWGEFA$ | 0.041 | -0.016 | - | **-0.163** | 0.003 |
| $MLC$ | **0.174** | **0.165** | **0.163** | - | **0.164** |
| $MLC_G$ | 0.055 | -0.029 | -0.003 | **-0.164** | - |

Table S8: Spearman correlation of the % difference in performance between the method in each row and the one each column with term path length to the ontology root. Statistically significant values (FDR $<$ 0.05) are shown in bold.

| | $PCC$ | $PCC + MR$ | $GAAWGEFA$ | $MLC$ | $MLC_G$ |
|---|---|---|---|---|---|
| $PCC$ | - | -0.007 | -0.035 | **-0.259** | -0.035 |
| $PCC + MR$ | 0.007 | - | 0.022 | **-0.246** | 0.015 |
| $GAAWGEFA$ | 0.035 | -0.022 | - | **-0.244** | -0.018 |
| $MLC$ | **0.259** | **0.246** | **0.244** | - | **0.255** |
| $MLC_G$ | 0.035 | -0.015 | 0.018 | **-0.255** | - |

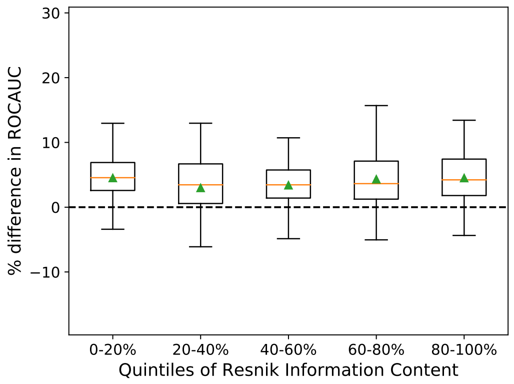

(a) *MR - PCC*

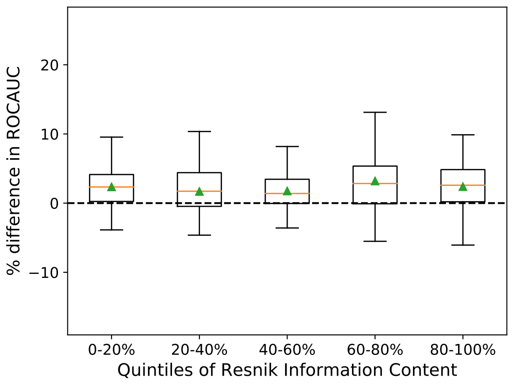

(b) *GAAWGEFA - PCC*

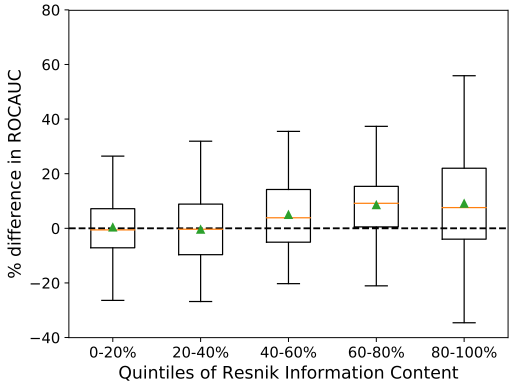

(c) *MLC - PCC*

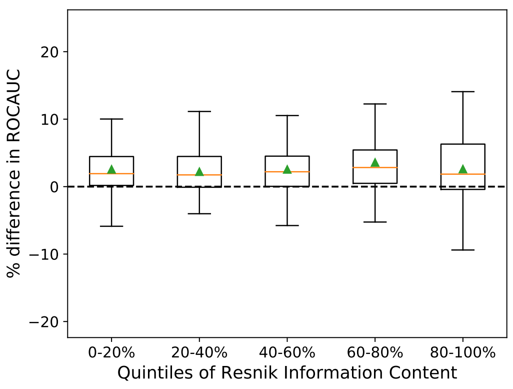

(d) $MLC_G$ - *PCC*

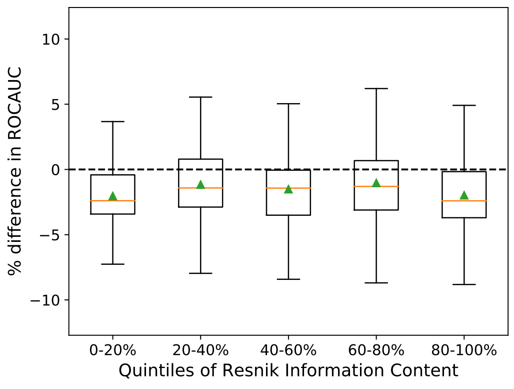

(e) *GAAWGEFA - MR*

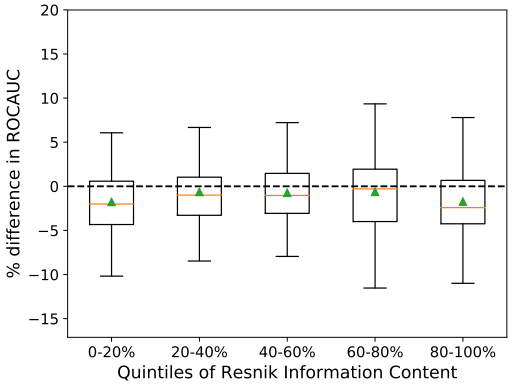

(f) $MLC_G$ - *MR*

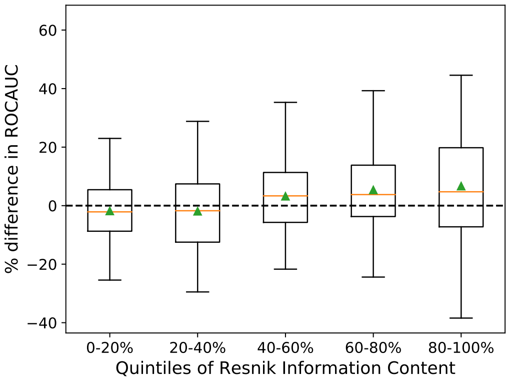

(g) *MLC - GAAWGEFA*

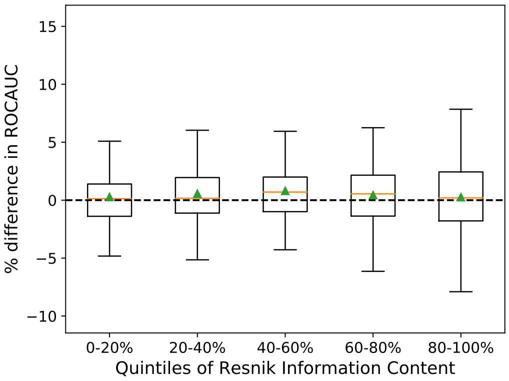

(h) $MLC_G$ - *GAAWGEFA*

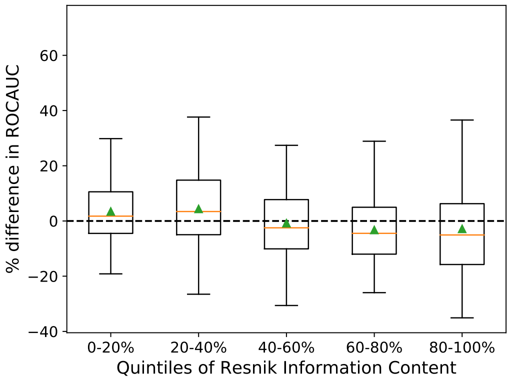

(i) $MLC_G$ - *MLC*

Figure S2: Percent difference in $ROCAUC$ of all pairs of methods as a function of Resnik Information Content. For each set of terms in each quintile of Information Content, the corresponding box includes the two middle quartiles of the percent difference for these terms. An orange line denotes the median and a green triangle the mean. The error bars extend to 1.5 times the range of the two middle quartiles and outlier points are shown as dots.

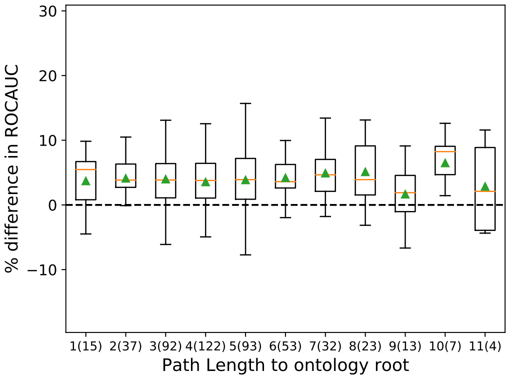

(a) *MR - PCC*

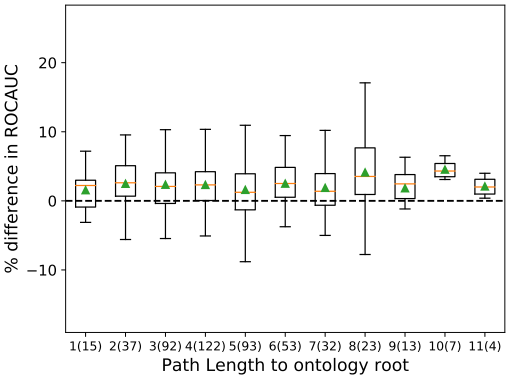

(b) *GAAWGEFA - PCC*

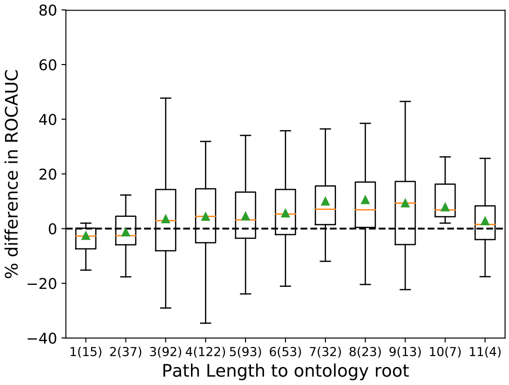

(c) *MLC - PCC*

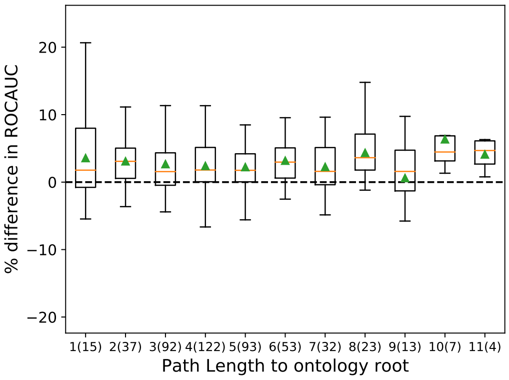

(d) $MLC_G$ - *PCC*

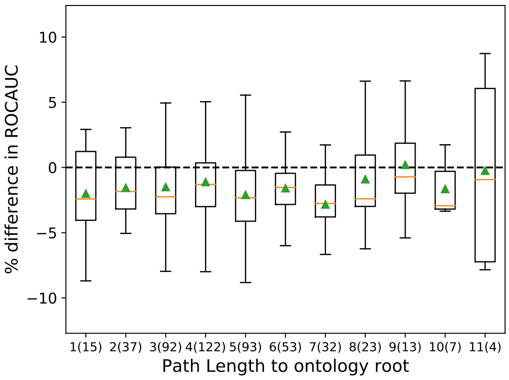

(e) *GAAWGEFA - MR*

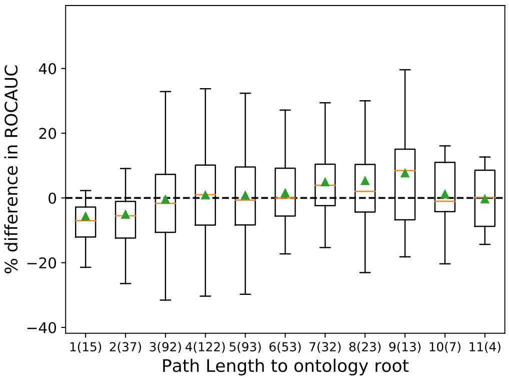

(f) *MLC - MR*

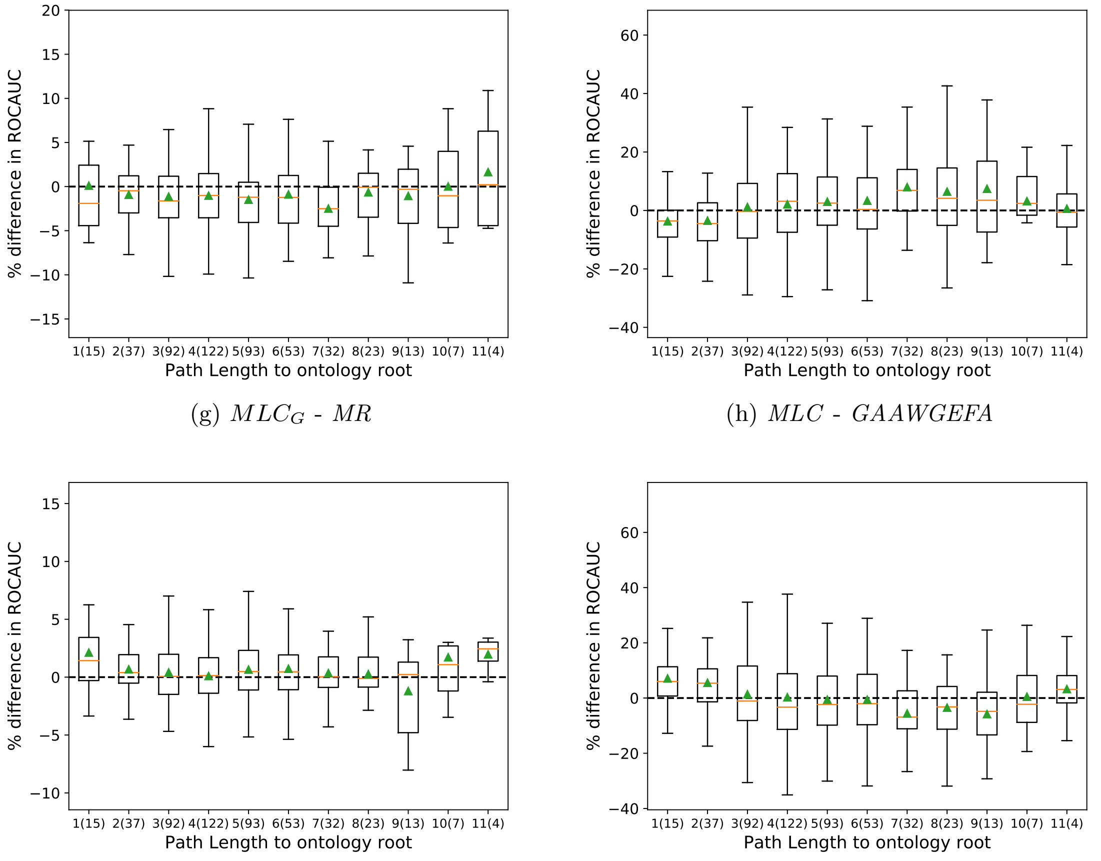

(g) $MLC_G$ - $MR$

(h) $MLC$ - $GAAWGEFA$

(i) $MLC_G$ - $GAAWGEFA$

(j) $MLC_G$ - $MLC$

Figure S3: Percent difference in $ROCAUC$ of all pairs of methods as a function of path length to the ontology root. For each set of terms with a given path length, the corresponding box includes the two middle quartiles of the percent difference for these terms. An orange line denotes the median and a green triangle the mean. The error bars extend to 1.5 times the range of the two middle quartiles and outlier points are shown as dots. The numbers in parentheses next to each path length level denote the number of terms in the dataset at that level.

### 7 Comparison of *GAAWGEFA* and *MLC* weights

In Figure S4 we show the distribution of the weight values of the "average" weight profile of *MLC*. To obtain that, we sort all term-specific weight profiles from the smallest to the largest value and then calculate the average weight value at each rank over all term profiles. Figure S4 also shows the weight distribution of the *GAAWGEFA*. We calculated the *PCC* between each term-specific profile and the *GAAWGEFA* profile and also between each term-specific profile and the $MLC_G$ profile. Figure S5 shows the distribution of these two similarities. In conclusion, the weights learned by *GAAWGEFA* were not correlated to the ones learned by *MLC* or $MLC_G$.

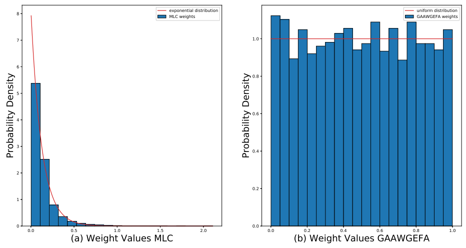

Figure S4: Histogram ($y$-axis) of the sample weight values ($x$-axis) for the average *MLC* profile (a) and the GAAWGEFA profile (b). The best fitting exponential distribution (a) and uniform distribution (b) are shown in red.

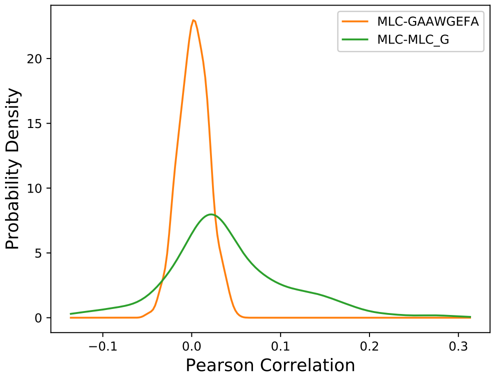

Figure S5: Distribution of the pairwise Pearson correlation values between the *MLC*-derived GO-term-specific weight profiles and the weight profile of $MLC_G$ (green) and *GAAWGEFA* (orange). The $x$ axis corresponds to the correlation values and the $y$ axis to the probability density.

### 8 Weight profile

Figure S6 shows the weight profile that *MLC* learned for term GO:1903047 (mitotic cell cycle process).

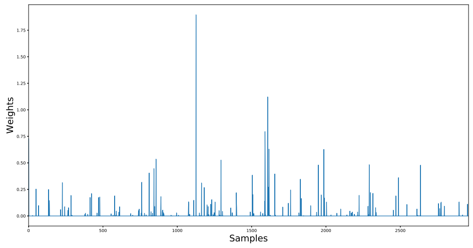

Figure S6: The weight profile learned by *MLC* for term GO:1903047. On the $x$ axis are the 2,959 RNA-Seq samples and on the $y$ axis the weight value of each sample.
